# Causes, temporal trends and the effects of urbanisation on admissions of wild raptors to rehabilitation centres in England and Wales

**DOI:** 10.1101/2022.02.10.479874

**Authors:** Connor T. Panter, Simon Allen, Nikki Backhouse, Elizabeth Mullineaux, Carole-Ann Rose, Arjun Amar

**Affiliations:** School of Geography, University of Nottingham, Nottingham, NG7 2RD, UK; Gower Bird Hospital, Sandy Lane, Southgate, Swansea, SA3 2EW, UK; Cuan Wildlife Rescue, The Signals, Stretton Road, Much Wenlock, Shropshire, TF13 6DD, UK; Secret World Wildlife Rescue, New Road, East Huntspill, Highbridge, Somerset, TA9 3PZ, UK; Wild Wings Birds of Prey, New Hall Lane, Risley, Croft, Warrington, Cheshire, WA3 6BH, UK; Fitzpatrick Institute of African Ornithology, DSI-NRF Centre of Excellence, University of Cape Town, Rondebosch, South Africa

**Keywords:** Birds of prey, conservation, mortality, morbidity, threats, wildlife rescue centres, rehabilitation

## Abstract

Data from wildlife rehabilitation centres can provide on-the-ground records of causes of raptor morbidity and mortality, allowing threat patterns to be explored throughout time and space. We provide an overview of native raptor admissions to four wildlife rehabilitation centres (WRCs) in England and Wales, quantifying the main causes of morbidity and mortality, trends over time and whether certain causes were more common in more urbanised areas between 2001-2019. Throughout the study period 14 raptor species were admitted totalling 3305 admission records. The Common Buzzard (*Buteo buteo;* 31%) and Tawny Owl (*Strix aluco*; 29%) were most numerous. Relative to the proportion of breeding individuals in Britain & Ireland, Peregrine Falcons (*Falco peregrinus*), Little Owls (*Athene noctua*) and Western Barn Owls (*Tyto alba*) were over-represented in the admissions data by 103%, 73% and 69%, respectively. Contrastingly Northern Long-eared Owls (*Asio otus*), Western Marsh Harriers (*Circus aeruginosus*) and Merlin (*Falco columbarius*) were under-represented by 187%, 163% and 126%, respectively. Across all species, vehicle collisions were the most frequent anthropogenic admission cause (22%) and orphaned young birds (10%) were most frequent natural admission cause. Mortality rate was highest for infection/parasite admissions (90%), whereas orphaned birds experienced lowest mortality rates (16%). For one WRC, there was a notable decline in admissions over the study period. Red Kite (*Milvus milvus*) admissions increased over time, whereas Common Buzzard and Common Kestrel admissions declined. There were significant declines in the relative proportion of persecution and metabolic admissions, and an increase in orphaned young birds. Urban areas were positively associated with persecution, building collisions and unknown trauma admissions, whereas vehicle collisions were associated with more rural areas. Many threats persist for raptors in England and Wales, however, have not changed substantially over the past two decades. Threats associated with urban areas, such as building collisions, may increase over time in line with human population growth and subsequent urban expansion.

## INTRODUCTION

Diurnal and nocturnal raptors are frequently used as ecological indicators due to their high positions within trophic networks (Buechley *et al*. 2019). Raptor species face a number of threats from anthropogenic activities such as direct and indirect poisoning (Hughes *et al*. 2013; Garvin *et al*. 2020), electrocution on powerlines (Lehman *et al*. 2007), road collisions (Gagné *et al*. 2015) and human persecution (Smart *et al*. 2010; Murgatroyd *et al*. 2019, Panter *et al*. 2021). For effective conservation programmes, the key detrimental impacts of anthropogenic activities need to be identified and evidenced-based conservation measures implemented to alleviate these threats (Holmes *et al*. 1993; Richardson & Miller 1997; Hernandez *et al*. 2018).

Several methods have been applied to quantify the effects of anthropogenic activities on raptors. Such approaches include screening for organic pollutants and contaminants (López *et al*. 2001; Chen *et al*. 2010), monitoring the dynamics of the illicit wildlife trade (Panter & White 2020), analysis of powerline collisions data (Bevanger 1998; Kolnegari *et al*. 2020), monitoring via remote tracking devices (Kendall & Virani 2012; McIntyre 2012; Panter *et al*. 2020, 2021) and wildlife rehabilitation admissions data (see Fix & Barrows 1990; Morishita *et al*. 1998; Wendell *et al*. 2002; Komnenou *et al*. 2005; Rodríguez *et al*. 2010; Molina-López *et al*. 2011; Molina-López & Darwich 2011; Thompson *et al*. 2013; Al Zoubi *et al*. 2020).

Raptor data from wildlife rehabilitation centres provide on-the-ground records of causes of morbidity and mortality and have been used to evaluate the health status of wild populations (Morishita *et al*. 1998; Wendell *et al*. 2002) and to explore trends in anthropogenic threats over time (Molina-López *et al*. 2011; Thompson *et al*. 2013). Rehabilitation and subsequent release of individuals back into the wild can help to buffer the negative effects of anthropogenic activities, especially for species of conservation concern (Mullineaux 2014; Montesdeoca *et al*. 2017a; Hernandez *et al*. 2018; Romero *et al*. 2019; Thomson *et al*. 2020; Dessalvi *et al*. 2021).

While several previous studies have explored morbidity and mortality of raptors based on admissions data to rehabilitations centres, most of these were based on data from a single centre, limiting their ability to explore patterns in admission causes over larger spatial scales. To our knowledge no studies have attempted to explore whether causes of morbidity or mortality differ depending on environmental features and very few have been conducted in the United Kingdom. For example, Kelly & Bland (2006) analysed admissions, diagnoses and outcomes of raptors admitted to a centre in England, focusing on a single species – the Eurasian Sparrowhawk *(Accipiter nisus).*

In this study, we compile and analyse raptor admissions data from four wildlife rehabilitation centres in western/south-western England and Wales. Firstly, we provide an overview of raptor admissions over a 19-year period (2001-2019), quantifying the most frequently admitted species and the main causes. We then explore whether numbers of commonly admitted species, and the types (anthropogenic vs natural) or causes of admission have changed over time for one rehabilitation centre, for which we had the longest run of data. Over the study period, urban cover in England and Wales has increased (Office for National Statistics 2021). Therefore, we predict an increase in anthropogenic admissions as a result of increasing human population growth and urban expansion over time (Seto *et al*. 2012). Certain threats may also have changed over time, for example over the study period the number of vehicles in England and Wales has increased (Department of Transport 2020), and subsequent raptor-vehicle collisions may also have increased over time. Finally, we expect that causes of admission will vary depending on the level of urbanisation. For example, we might expect that urbanisation increases the probability of admissions due to building or vehicle collisions in line with previous findings (Loss *et al*. 2014; Garcês *et al*. 2020).

Therefore, we explore whether the level of urbanisation (where the individual birds were found) is associated with higher probabilities of certain admission causes.

## METHODS

### Study area

We collated admission records of native raptors admitted to wildlife rehabilitation centres (WRC) located within a study area totalling *c*. 46,000 km^2^ in south-western Britain (Fig. 1). The landscape within our study area is dominated by agriculture but also includes the major cities of Greater Manchester, Birmingham, Bristol and Cardiff which have populations of *c*. 2.8 million, 2.6 million, 690,000 and 495,000 people, respectively (United Nations 2014). Our study area also includes the Brecon Beacons National Park, seven ‘Areas of Outstanding Natural Beauty’ (AONB) and numerous ‘Sites of Special Scientific Interest’ (SSSI) including the West Pennine Moors, Wyre Forest and the Quantock Hills.

**Figure 1.**
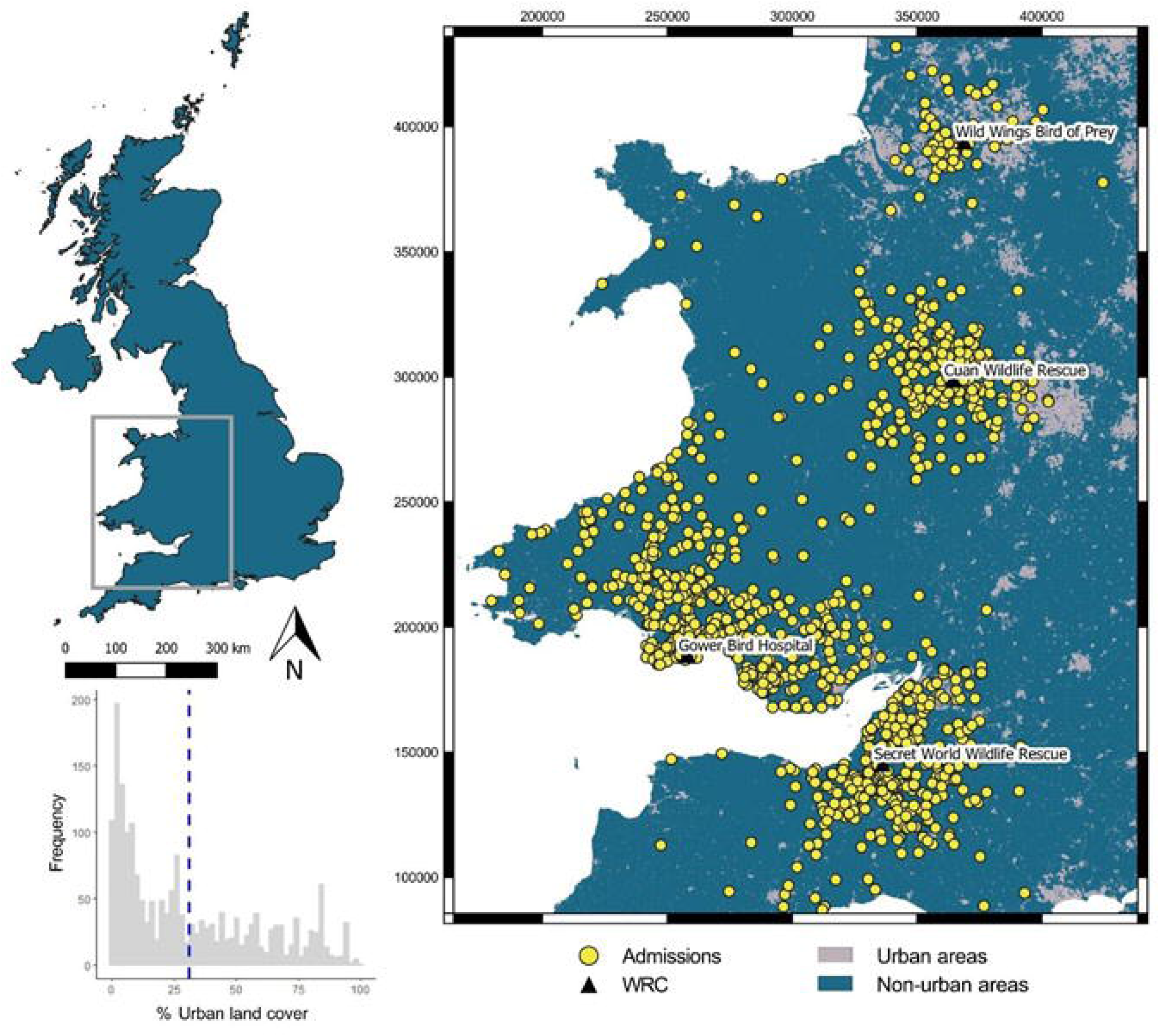
Spatial distribution for 14 species of diurnal and nocturnal raptors admitted to four wildlife rehabilitation centres (WRC) between 2001-2019 in England and Wales. Geo-referenced admissions with 2 km buffers (N = 1915) shown in relation to urban land cover. Histogram shows the frequency of urban land cover scores within each 2 km buffer and the mean (31%) denoted by the blue dashed line. Map Coordinate Reference System: EPSG 27700 British National Grid.

### Data collection

Wildlife rehabilitation centres were invited to participate in the study via email correspondence. Four WRC supplied data on raptor admissions to their centres: Cuan Wildlife Rescue (lat/long: 52.590, −2.573), Gower Bird Hospital (51.580, −4.099), Secret World Wildlife Rescue (51.206, −2.964) and Wild Wings Birds of Prey (53.444, −2.522). From their admissions records the following data were collected for each individual admitted: 1) species, 2) sex (male/female), 3) age (juvenile/adult; <1 calendar year/>1cy), 4) admission date, 5) cause of admission, 6) location of incident (at the finest spatial scale available) and 7) outcome (deceased/released/kept in captivity). These data spanned a 19-year period from 21^st^ January 2001 to 26^th^ December 2019.

### Classifying causes of morbidity and mortality

To increase comparability with other studies, classification of admission causes followed categories previously defined by existing studies (see Molina-López *et al*. 2011; Molina-López & Darwich 2011). Upon admission, birds were examined by trained wildlife carers and the admission notes associated with each record were used to assign each admission to the following ‘types’ (‘ANTHROPOGENIC’, ‘NATURAL’ and ‘UNKNOWN’) and more detailed ‘causes’ (see supplementary Appendix A for an overview of all admission types, causes, codes and pooled miscellaneous causes). When causes could not be ascertained, admission type was categorised as ‘UNKNOWN’ which, included the causes: ‘undetermined’ (reason unknown and no injury to bird) and ‘unknown trauma’ (reason unknown but the bird was physically injured).

### Landscape and demographic variables

To explore urbanisation effects on types and causes of raptor admissions, we used only the geo-referenced admissions (N = 1915). For these, we extracted land cover data and calculated the proportion of urban habitat within a 2 km buffer. Land cover data was downloaded on 30^th^ April 2020 from the EDINA Environment Digimap Service (Land Cover Map 2015; https://digimap.edina.ac.uk/). Land cover data was derived from the ‘LCM2015’ dataset in raster format at 25 m resolution, which closely aligned with the timescale of the majority of the admissions. All spatial data extraction were performed in QGIS 3.12.3 with the GRASS 7.4.1 extension (QGIS Development Team 2019). We reclassified the land cover data using the *r.reclass* function and a new binary raster layer was created (1 = ‘urban’ + ‘suburban’ and 0 = all other land cover types). Summary statistics were then computed using the base function *Zonal Statistics* to calculate percentage urban cover within each 2 km buffer.

### Statistical analysis

All statistical analyses were performed in R version 3.6.3 (R Core Team, 2020). Data were analysed used Generalized Linear Models (GLMs) with either binomial (for binary models) or Poisson (for count data) distributions, and the respective conical link functions (See Appendix B for list of models). For binomial data, we fitted a two-vector response variable using the cbind function. For Poisson GLMs where overdispersion was detected we fitted the models with a quasi-Poisson distribution.

We explored mortality (binary: 1 = bird died or was euthanised termed ‘deceased’ and 0 = bird released or kept captive termed ‘not deceased’) as a response variable, with explanatory variables of either admission type or cause. We explore trends over time using only data from Gower Bird Hospital as it was the only WRC with the longest run of data. Using these data, we fitted year as the explanatory variable and fitted a series of separate GLMs with the following response variables: 1) total count of admission each year, irrespective of cause and including unknown causes (Poisson model). 2) Total count of admission each year for the seven most frequently admitted species (with ≥ 30 admissions). 3) Relative proportion, per year, of admission causes (with ≥ 30 admissions). 4) admission type, anthropogenic or natural (binomial model).

The effects of urbanisation on types and causes of admissions were explored using a series of GLMMs in the package ‘lme4’ (Bates *et al*. 2015). For each admission, a binary metric was created (1 = matching admission type and 0 = no match) for each admission type (i.e., anthropogenic, natural or unknown), or admission cause (where there were ≥ 30 admissions, i.e., vehicle collisions, trauma, undetermined, orphaned, building collisions, metabolic, infections/parasites and persecution). These models were then run with ‘binary admission type/cause’ fitted as the response term and ‘% urban land cover’ fitted as the explanatory term. We used binomial error distributions and ‘logit’ link functions with ‘centrelD’ included as a random term to control for the lack of independence between admissions from the same centre (Appendix B).

We examined whether certain species were over- and under-represented within our admissions data by calculating the percentage difference between the relative proportion of breeding individuals in Britain and Ireland, and the proportion of admitted individuals, per species, to each WRC. Breeding population data were derived from the British Trust for Ornithology’s BirdFacts database (Robinson 2005; https://www.bto.org/understanding-birds/birdfacts).

## RESULTS

Across the 19-year study period, we recorded a total of 3305 admissions, comprising of 14 species, (Table 1), with 1919 (58%) of admissions being diurnal species and 1386 (42%) being nocturnal species. The diurnal raptors comprised of nine species, the Common Buzzard *(Buteo buteo)* (N =1035; 31%) being the most frequently admitted species, followed by the Eurasian Sparrowhawk *(Accipiter nisus)* (N = 457; 14%) and then the Common Kestrel *(Falco tinnunculus)* (N = 269; 8%). The Tawny Owl *(Strix aluco)* (N = 967; 29%) was the second most frequently admitted of all species and the most frequently admitted nocturnal species, followed by the Western Barn Owl *(Tyto alba)* (N = 283; 9%) and the Little Owl (*Athene noctua*) (N = 118; 4%).

**Table 1.** Demographics of diurnal and nocturnal raptor species admitted to four wildlife rehabilitation centres in England and Wales between 2001-2019. Demographic proportions calculated per species, totals calculated based on total number of admissions. *Proportions calculated using total diurnal and nocturnal values.

| Species | Sex |  |  | Age |  |  | Total (%) |
| --- | --- | --- | --- | --- | --- | --- | --- |
|  | Male<br>(%/sp.) | Female<br>(%/sp.) | Unknown<br>(%/sp.) | Adult<br>(%/sp.) | Juvenile<br>(%/sp.) | Unknown<br>(%/sp.) |  |
| *Diurnal* |  |  |  |  |  |  |  |
| Common Buzzard (*Buteo buteo*) | 107 (10) | 120 (12) | 808 (78) | 615 (59) | 287 (28) | 133 (13) | 1035 (31) |
| Eurasian Sparrowhawk (*Accipiter nisus*) | 92 (20) | 129 (28) | 236 (52) | 240 (53) | 158 (35) | 59 (13) | 457 (14) |
| Common Kestrel (*Falco tinnunculus*) | 48 (18) | 38 (14) | 183 (68) | 114 (42) | 122 (45) | 33 (12) | 269 (8) |
| Peregrine Falcon (*Falco peregrinus*) | 28 (33) | 22 (26) | 34 (40) | 44 (52) | 36 (43) | 4 (5) | 84 (3) |
| Red Kite (*Milvus milvus*) | 3 (8) | 4 (11) | 29 (81) | 27 (75) | 9 (25) |  | 36 (1) |
| Eurasian Hobby (*Falco subbuteo*) | 1 (6) | 2 (12) | 14 (82) | 12 (71) | 1 (6) | 4 (24) | 17 (1) |
| Northern Goshawk (*Accipiter gentilis*) | 5 (31) | 6 (38) | 5 (31) | 3 (19) | 13 (81) |  | 16 (<1) |
| Merlin (*Falco columbarius*) | 1 (25) | 1 (25) | 2 (50) | 1 (25) | 3 (75) |  | 4 (<1) |
| Western Marsh Harrier (*Circus aeruginosus*) |  |  | 1 (100) |  |  | 1 (100) | 1 (<1) |
| Total diurnal* | 285 (15) | 322 (17) | 1312 (68) | 1056 (55) | 629 (33) | 234 (12) | 1919 (58) |
| *Noctural* |  |  |  |  |  |  |  |
| Tawny Owl (*Strix aluco*) | 35 (4) | 21 (2) | 911 (94) | 474 (49) | 359 (37) | 134 (14) | 967 (29) |
| Western Barn Owl (*Tyto alba*) | 39 (14) | 54 (19) | 190 (67) | 156 (55) | 98 (35) | 29 (10) | 283 (9) |
| Little Owl (*Athene noctua*) | 1 (1) | 1 (1) | 116 (98) | 41 (35) | 63 (53) | 14 (12) | 118 (4) |
| Short-eared Owl (*Asio flammeus*) |  | 3 (19) | 13 (81) | 12 (75) | 3 (19) | 1 (6) | 16 (<1) |
| Northern Long-eared Owl (*Asio otus*) |  |  | 2 (100) | 2 (100) |  |  | 2 (<1) |
| Total nocturnal* | 75 (5) | 79 (6) | 1232 (89) | 685 (49) | 523 (38) | 178 (13) | 1386 (42) |
| Total admissions | 360 (11) | 401 (12) | 2544 (77) | 1741 (53) | 1152 (35) | 412 (12) | 3305 (100) |

Only 761 (23%) admitted birds were successfully sexed, of these 47% were males and 53% were females. Age was determined for 2893 (88%) admissions with adults (>1cy) representing 60% and juveniles (<1cy) 40% of these aged individuals (Table 1).

### Admission types and causes

Unknown admission types were the most numerous comprising nearly half of all admissions (n=1510; 46%), followed by anthropogenic (n=1215; 37%) then natural causes (n=580; 17%; Table 2). Classifying admissions by the more detailed ‘causes’ revealed 855 (26%) of all admissions were associated with ‘unknown trauma’ (Table 2). The most frequent anthropogenic admission cause was ‘vehicle collisions’ (n=732; 22% of all admissions; 60% of anthropogenic admissions). For natural admissions, orphaned young birds was the most frequent cause (n= 315; 10% of all admissions, 54% of natural admissions; Table 2).

**Table 2.** Admission types and causes for 14 species of diurnal and nocturnal raptors, admitted to four wildlife rehabilitation centres in England and Wales between 2001-2019. *Proportions calculated using total diurnal and nocturnal values. Causes: ‘attack’ = attacked by pet, ‘build’ = building collisions, ‘elec’ = electrocutions, ‘fence’ = fencing/entanglements, ‘habitat’ = habitat destruction, ‘pers’ = persecutions, ‘veh’ = vehicle collisions, ‘infect’ = infection/parasites, ‘metab’ = metabolic, ‘orph’ = orphaned, ‘pred’ = predation, ‘trauma’ = unknown trauma and ‘undet’ = undetermined. See Table S1 for full cause descriptions.

| Species admitted | Anthropogenic (%/sp.) |  |  |  |  |  |  | Natural (%/sp.) |  |  |  | Unknown (%sp.) |  | Total (%) |
| --- | --- | --- | --- | --- | --- | --- | --- | --- | --- | --- | --- | --- | --- | --- |
|  | attack | build | elec | fence | habitat | pers | veh | infect | metab | orph | pred | trauma | undet |  |
| *Diurnal* |  |  |  |  |  |  |  |  |  |  |  |  |  |  |
| Common Buzzard<br>(*Buteo buteo*) |  | 26 (3) | 10 (1) | 12 (1) | 3 (<1) | 25 (2) | 262 (25) | 30 (3) | 68 (7) | 20 (2) | 8 (1) | 333 (32) | 238 (23) | 1035 (31) |
| Eurasian Sparrowhawk<br>(*Accipiter nisus*) | 19 (4) | 105 (23) | 1 (<1) | 9 (2) |  | 15 (3) | 40 (9) | 13 (3) | 8 (2) | 6 (1) | 6 (1) | 163 (36) | 72 (16) | 457 (14) |
| Common Kestrel<br>(*Falco tinnunculus*) | 1 (<1) | 15 (6) | 2 (1) | 2 (1) | 1 (<1) | 1 (<1) | 37 (14) | 3 (1) | 26 (10) | 38 (14) |  | 80 (30) | 63 (23) | 269 (8) |
| Peregrine Falcon<br>(*Falco peregrinus*) |  | 2 (2) |  | 3 (4) |  | 5 (6) | 6 (7) | 1 (1) | 2 (2) | 10 (12) | 2 (2) | 38 (45) | 15 (18) | 84 (3) |
| Red Kite (*Milvus milvus*) |  | 5 (14) |  |  |  | 1 (3) | 9 (25) |  |  | 2 (6) |  | 4 (11) | 15 (42) | 36 (1) |
| Eurasian Hobby<br>(*Falco subbuteo*) |  | 3 (18) |  |  |  |  | 4 (24) |  |  |  | 1 (6) | 5 (29) | 4 (24) | 17 (1) |
| Northern Goshawk<br>(*Accipiter gentilis*) |  | 3 (19) | 1 (6) |  |  | 1 (6) |  |  | 1 (6) | 1 (6) |  | 7 (44) | 2 (13) | 16 (<1) |
| Merlin (*Falco columbarius*) |  | 1 (25) |  |  |  |  |  |  |  |  |  | 3 (75) |  | 4 (<1) |
| Western Marsh Harrier (*Circus aeruginosus*) |  |  |  |  |  |  |  |  |  |  |  | 1 (100) |  | 1 (<1) |
| Total diurnal* | 20 (1) | 160 (8) | 14 (1) | 26 (1) | 4 (<1) | 48 (3) | 358 (19) | 47 (2) | 105 (5) | 77 (4) | 17 (1) | 634 (33) | 409 (21) | 1919 (58) |
| *Nocturnal* |  |  |  |  |  |  |  |  |  |  |  |  |  |  |
| Tawny Owl (*Strix aluco*) | 8 (1) | 82 (8) |  | 41 (4) | 8 (1) | 17 (2) | 290 (30) | 35 (4) | 24 (2) | 154 (16) | 6 (1) | 133 (14) | 169 (17) | 967 (29) |
| Western Barn Owl<br>(*Tyto alba*) | 3 (1) | 9 (3) | 1 (<1) | 2 (1) | 6 (2) | 7 (2) | 66 (23) | 6 (2) | 11 (4) | 50 (18) | 4 (1) | 60 (21) | 58 (20) | 283 (9) |
| Little Owl (*Athene noctua*) | 5 (4) | 13 (11) |  |  | 5 (4) | 1 (1) | 17 (14) |  | 5 (4) | 34 (29) | 5 (4) | 18 (15) | 15 (13) | 118 (4) |
| Short-eared Owl<br>(*Asio flammeus*) |  |  |  |  | 1 (6) | 1 (6) | 1 (6) |  |  |  |  | 9 (56) | 4 (25) | 16 (<1) |
| Northern Long-eared Owl (*Asio otus*) |  |  |  |  |  | 1 (50) |  |  |  |  |  | 1 (50) |  | 2 (<1) |
| Total nocturnal* | 16 (1) | 104 (8) | 1 (<1) | 43 (3) | 20 (1) | 27 (2) | 374 (27) | 41 (3) | 40 (3) | 238 (17) | 15 (1) | 221 (16) | 246 (18) | 1386 (42) |
| Total | 36 (1) | 264 (8) | 15 (<1) | 69 (2) | 24 (<1) | 75 (2) | 732 (22) | 88 (3) | 145 (4) | 315 (10) | 32 (1) | 855 (26) | 655 (20) | 3305 (100) |

When exploring only identified admission causes (excluding all unknown admission causes), vehicle collisions were the most common cause for five species including the Common Buzzard (56%; N = 262/464), Red Kite *(Milvus milvus;* 53%; 9/17), Eurasian Hobby *(Falco subbuteo;* 50%; 4/8), Tawny Owl (44%; 290/665) and Western Barn Owl (40%; 66/165) (Table 2). For the two most admitted diurnal species, the Common Buzzard and Eurasian Sparrowhawk, unknown trauma was the most common admission cause (Fig. 2). Main admission causes for Tawny Owls were vehicle collisions and orphaned young birds, comprising 40% and 49% of admissions, respectively (Table 2; Fig. 2).

**Figure 2.**
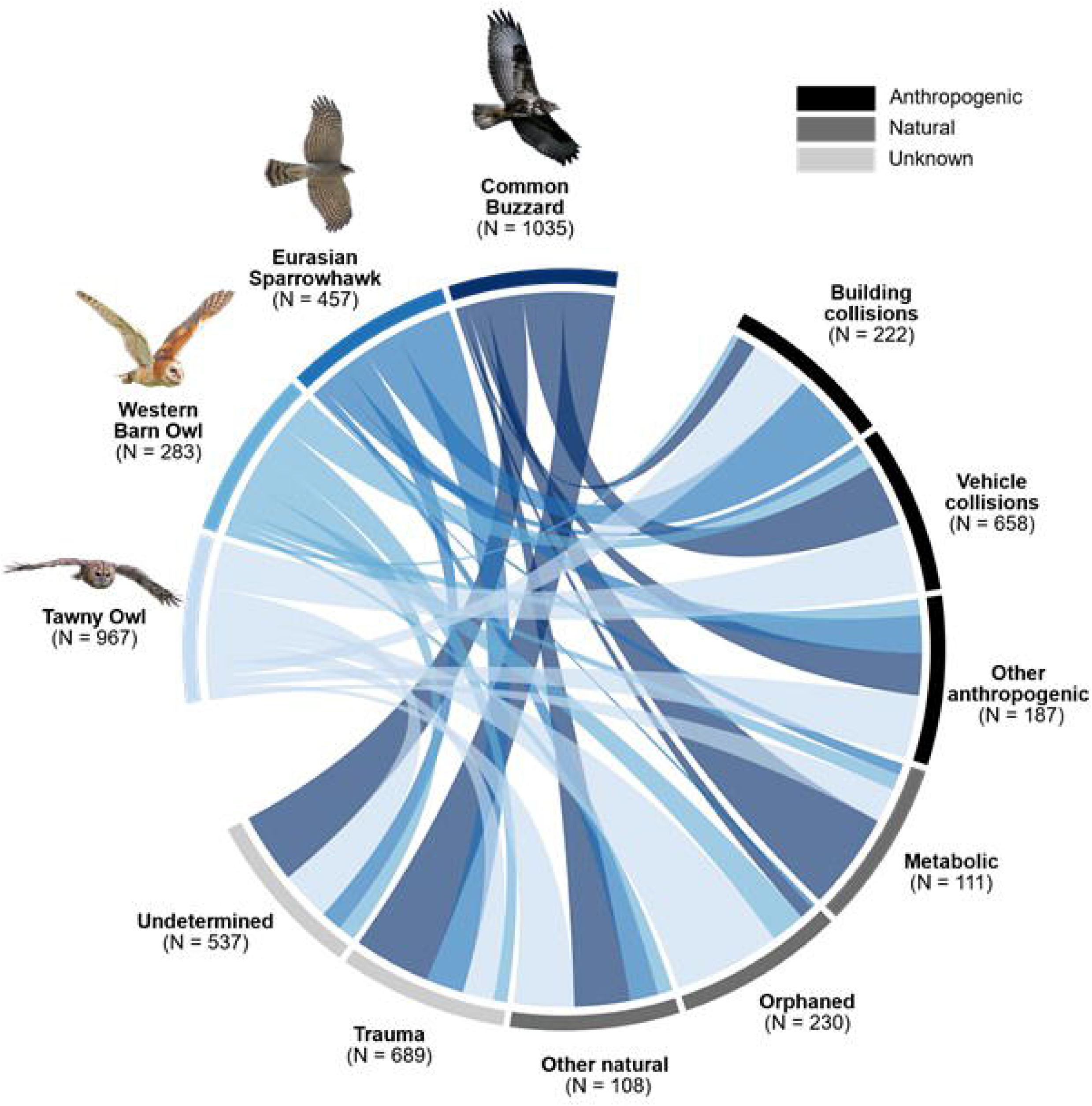
Admission causes for the top two most common diurnal and nocturnal raptor species admitted to four wildlife rehabilitation centres between 2001-2019 (N = 3011). Only the two most common admission causes per type (anthropogenic, natural and unknown) shown, other causes pooled into respective categories: ‘Other anthropogenic’ causes include ‘attacked’ (N = 30), ‘fencing/entanglement’ (N = 64), ‘electrocution’ (N = 12), ‘Habitat destruction’ (N = 17) and ‘Persecution’ (N = 64). ‘Other natural’ causes include ‘Infection/parasites’ (N = 84) and ‘Predation’ (N = 24).

Juvenile birds were approximately four times more likely to be admitted due to natural admissions than adults (430 vs. 112 admissions, respectively), and one and half times more likely to be admitted due to metabolic causes, e.g., emaciation or starvation, (79 vs. 54 admissions, respectively). Orphaned young birds totalled 10% (315) of all admissions and were the most frequent known admission cause for the Common Kestrel (14%; 38/269), Little Owl (29%; 34/118) and Peregrine Falcon *(Falco peregrinus;* 12%; 10/84) (Table 2).

### Outcome of admissions

From all admissions, 60% resulted in the death or euthanasia of the bird, 39% resulted in the release of the bird and just 1% of birds were kept in captivity post-admission (Table 3). Those admitted for anthropogenic reasons had a significantly higher mortality rate (57%) than those admitted for natural reasons (40%) *(z* = 6.483, *P* < 0.0001) (Fig 3a; Table 3; Appendix C). Mortality probabilities differed among the most common admission causes (Fig. 3b). Raptors admitted due to infection/parasites had a substantially higher mortality rate (90%) compared to other known admission causes, whereas orphaned birds had a significantly lower mortality rate (16%) than other known admission causes (Fig. 3b; Table 3; Appendix C).

**Table 3.** Overview of admission type, causes and outcomes for all raptor admissions to four wildlife rehabilitation centres in England and Wales between 2001-2019. Outcome proportions calculated per admission cause, totals based on total number of admissions. *Proportions calculated using total admission type values.

| Type | Cause | Outcome |  |  | Total (%) |
| --- | --- | --- | --- | --- | --- |
|  |  | Kept captive<br>(%/cause) | Deceased/euthanized (%/cause) | Released<br>(%/cause) |  |
| Anthropogenic | Attacked by pet | 0 (0) | 23 (64) | 13 (36) | 36 (1) |
|  | Building collision | 1 (< 1) | 136 (52) | 127 (48) | 264 (8) |
|  | Electrocution | 0 (0) | 11 (73) | 4 (27) | 15 (<1) |
|  | Fencing/entanglement | 1 (2) | 33 (49) | 35 (51) | 69 (2) |
|  | Habitat destruction | 5 (21) | 3 (16) | 16 (67) | 24 (1) |
|  | Persecution | 1 (1) | 39 (53) | 35 (47) | 75 (2) |
|  | Vehicle collision | 1 (< 1) | 438 (60) | 293 (40) | 732 (22) |
| Total anthropogenic* |  | 9 (<1) | 683 (56) | 523 (43) | 1215 (37) |
| Natural | Infection/parasites | 1 (1) | 79 (91) | 8 (9) | 88 (3) |
|  | Metabolic | 0 (0) | 84 (58) | 61 (42) | 145 (4) |
|  | Orphaned | 25 (8) | 50 (17) | 240 (76) | 315 (10) |
|  | Predation | 0 (0) | 21 (66) | 11 (34) | 32 (1) |
| Total natural* |  | 26 (5) | 234 (40) | 320 (55) | 580 (18) |
| Unknown | Trauma | 3 (< 1) | 689 (81) | 163 (19) | 855 (26) |
|  | Undetermined | 5 (< 1) | 368 (57) | 282 (43) | 655 (20) |
| Total unknown* |  | 8 (1) | 1057 (70) | 445 (29) | 1510 (46) |
| Total admissions |  | 43 (1) | 1974 (60) | 1288 (39) | 3305 (100) |

**Figure 3.**
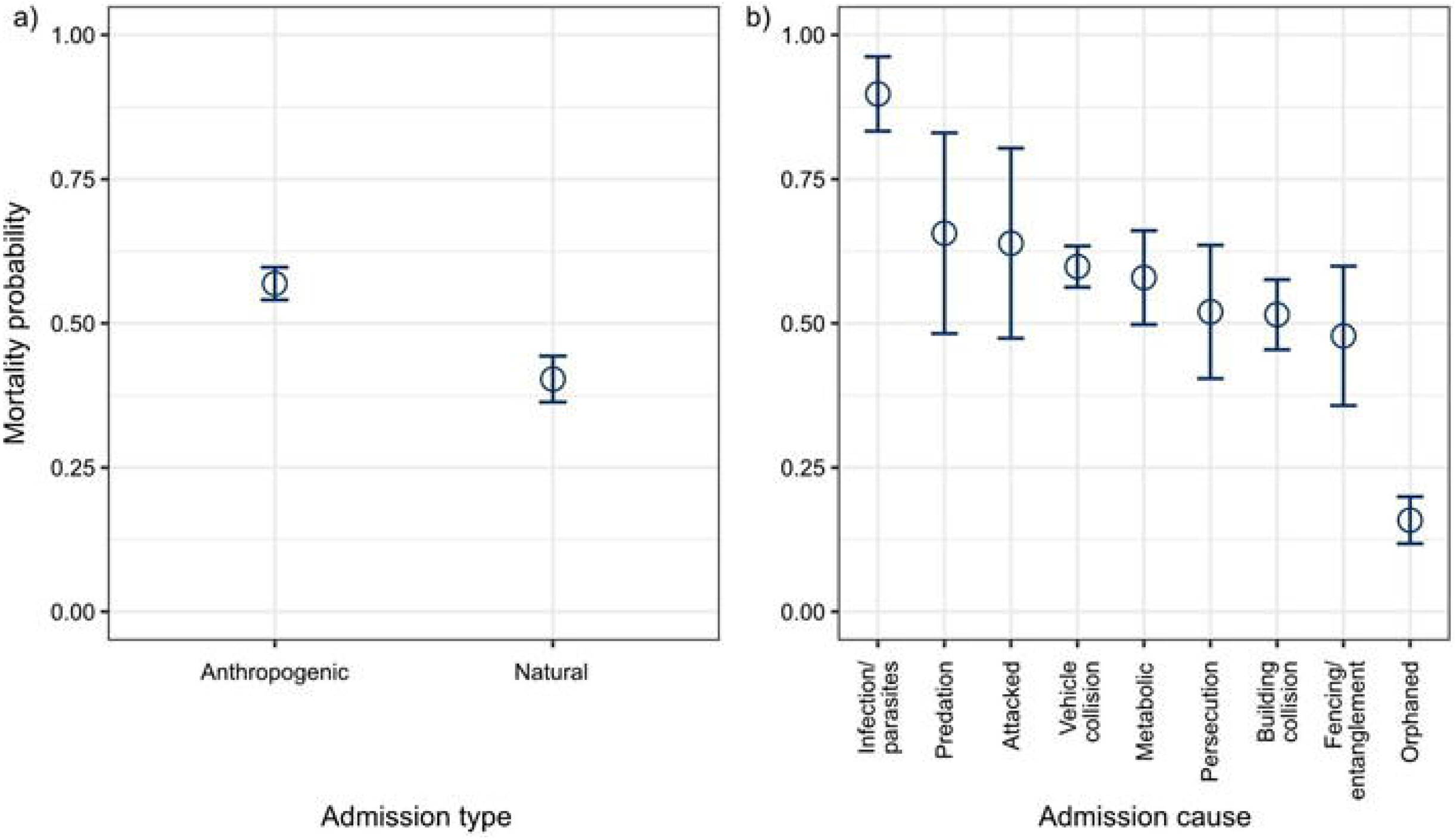
Differences in mortality probabilities for raptors admitted to four wildlife rehabilitation centres in England and Wales, between 2001-2019, in relation to identified a) admission types and b) admission causes. Data for ‘unknown’ admission type not shown. Error bars represent 95% confidence intervals.

### Trends over time in raptor admissions

Between 2001 and 2019, there was a notable decline in raptor admissions to Gower Bird Hospital when analysing all admission types (*t*_1,17_ value = −2.164, *P* < 0.05). However, the relative proportion of known anthropogenic vs. natural admissions admitted to Gower Bird Hospital did not change over time (*z*_1,17_ = −1.554, *P* = 0.120). Over this period, there was a significant increase in the number of Red Kites admitted (*t*_1,17_ = 4.703, *P* < 0.001) (Fig. 4). Conversely, there were significant declines in the number of Common Buzzards (*t*1,17 = −2.407, *P* < 0.05) and Common Kestrels admitted (*t*_1,17_ = −4.031, *P* < 0.001) (Fig. 4; Appendix D). We also saw a significant decline in the relative proportion of persecution and metabolic related admissions, and a significant increase in orphaned young birds, admitted to Gower Bird Hospital throughout the study period (Table 4).

**Figure 4.**
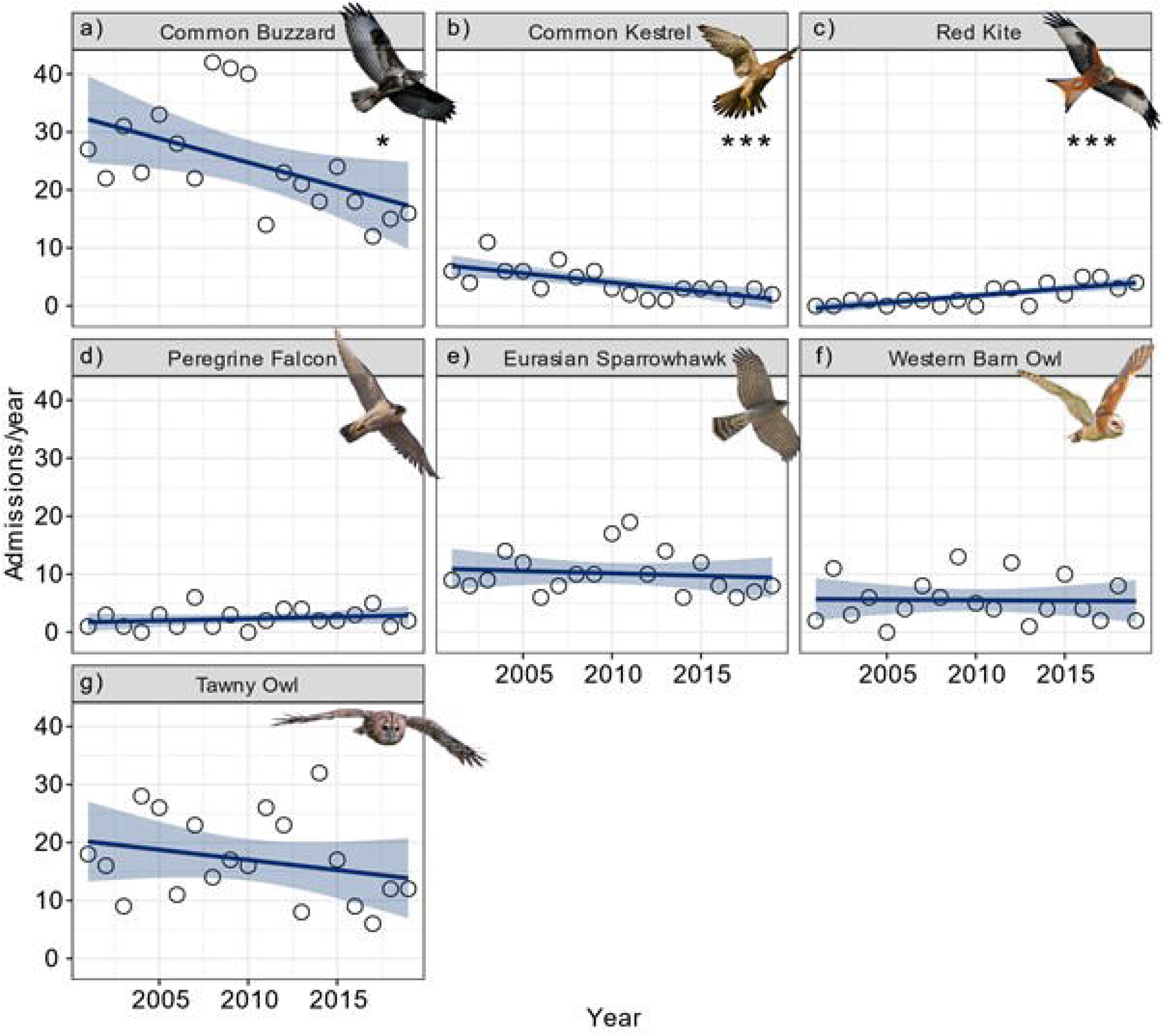
Trends over time for the seven most common raptor species admitted to Gower Bird Hospital between 2001-2019. a) Common Buzzard (*Buteo buteo*; N = 470), b) Common Kestrel *(Falco tinnunculus;* N = 77), c) Red Kite *(Milvus milvus;* N = 34), d) Peregrine Falcon *(Falco peregrinus;* N = 44), e) Eurasian Sparrowhawk *(Accipiter nisus;* N = 193), f) Western Barn Owl *(Tyto alba;* N = 105) and g) Tawny Owl *(Strix aluco;* N = 323). Significant trends over time denoted by ‘***’ = *P* < 0.001 and ‘*’ = *P* < 0.05.

**Table 4.** Trends over time in the relative proportion, per year, of admission causes for 1237 raptors admitted to Gower Bird Hospital between 2001-2019. Data analysed using a series of Generalized Linear Models fitted with quasi-Poisson error distributions to control for overdispersion. Only admission causes ≥ 30 were included. **Bold** = statistically significant causes. N = sample size, SE = standard error, df = degrees of freedom.

| Admission cause | N | Estimate ± SE | *t* | df | *P* |
| --- | --- | --- | --- | --- | --- |
| *Anthropogenic* |  |  |  |  |  |
| Building collision | 113 | -0.012 ± 0.020 | -0.607 | 18 | 0.552 |
| **Persecution** | **38** | **-0.074 ± 0.033** | **-2.258** | - | **< 0.05** |
| Vehicle collision | 322 | 0.008 ± 0.014 | 0.563 | - | 0.581 |
| *Natural* |  |  |  |  |  |
| Infection/parasites | 41 | 0.042 ± 0.039 | 1.081 | - | 0.295 |
| **Metabolic** | **78** | **-0.072 ± 0.033** | **-2.149** | - | **< 0.05** |
| **Orphaned** | **97** | **0.066 ± 0.028** | **2.302** | - | **< 0.05** |
| *Unknown* |  |  |  |  |  |
| Trauma | 312 | -0.011 ± 0.015 | -0.766 | - | 0.454 |
| Undetermined | 236 | 0.009 ± 0.020 | 0.466 | - | 0.647 |

### Effects of urbanisation

From 3305 admissions, 1915 (58%) were geo-referenced. For these geo-referenced admissions, the mean percentage urban land cover within the 2 km diameter buffers was 31 ± 28% (± SD) (Fig. 1). We found no significant association between the proportion of urbanisation for each geo-referenced admission and the probability that the admission was caused by anthropogenic (*z*_1,1914_ = 0.940, *P* = 0.347), natural (*z*_1,1914_ = −1.085, *P* = 0.278) or unknown factors (*z*_1,1914_ = −0.118, *P* = 0.906). We did, however, find a significant positive association between urbanisation and the probability of admission cause being building collisions, persecution or unknown trauma (Table 5). In the least urbanised areas, probability of admission being attributed to a building collision was only *c*. 7% but increased to *c*. 18% in the most urbanised areas. Likewise, persecution increased from *c*. 2.5% in the least urbanised areas to around 8% in the most urbanised areas. In contrast, vehicle collision admissions were negatively associated with urbanisation, with a considerably higher probability of admissions being attributed to vehicle collisions in less urbanised areas – this was also the case for undetermined admission causes (Table 5). Urbanisation was not associated with the probability of admission being attributed to any natural admission causes including infection/parasites, metabolic or orphaned young birds (Table 5).

**Table 5.** Effects of urbanisation on causes of admission for raptors admitted to four wildlife rehabilitation centres in England and Wales between 2001-2019. Data analyses using a series of Generalized Linear Mixed Models fitted with binomial error distributions and ‘logit’ link functions. **Bold** = statistically significant causes. Values computed using only geo-referenced admissions with 2 km diameter buffers.

| Admission cause | N | Estimate ± SE | *z* | df | *P* |
| --- | --- | --- | --- | --- | --- |
| *Anthropogenic* |  |  |  |  |  |
| **Building collision** | **136** | **0.011 ± 0.003** | **3.503** | **1109** | **< 0.001** |
| **Persecution** | **49** | **0.010 ± 0.005** | **2.047** | - | **< 0.05** |
| **Vehicle collision** | **503** | **-0.005 ± 0.002** | **-2.533** | - | **< 0.05** |
| *Natural* |  |  |  |  |  |
| Infection/parasites | 64 | -0.001 ± 0.005 | -0.223 | - | 0.824 |
| Metabolic | 105 | -0.005 ± 0.004 | -1.464 | - | 0.143 |
| Orphaned | 165 | 0.0005 ± 0.003 | 0.178 | - | 0.859 |
| *Unknown* |  |  |  |  |  |
| **Trauma** | **456** | **0.004 ± 0.002** | **1.980** | - | **< 0.05** |
| **Undetermined** | **349** | **-0.005 ± 0.002** | **-2.529** | - | **< 0.05** |

### Representation of raptor species

Compared to the relative proportion of breeding individuals in Britain and Ireland, some species were under- and over-represented within our admissions data (Fig. 5; Appendix E). For example, Peregrine Falcons, Little Owls and Western Barn Owls were over-represented in our admissions data by 103%, 73% and 69%, respectively (Fig. 5; Appendix E). Contrastingly, Northern Long-eared Owls (*Asio otus*), Western Marsh Harriers (*Circus aeruginosus*) and Merlin (*Falco columbarius*) were under-represented in our admissions data by 187%, 163% and 126%, respectively (Fig. 5; Appendix E).

**Figure 5.**
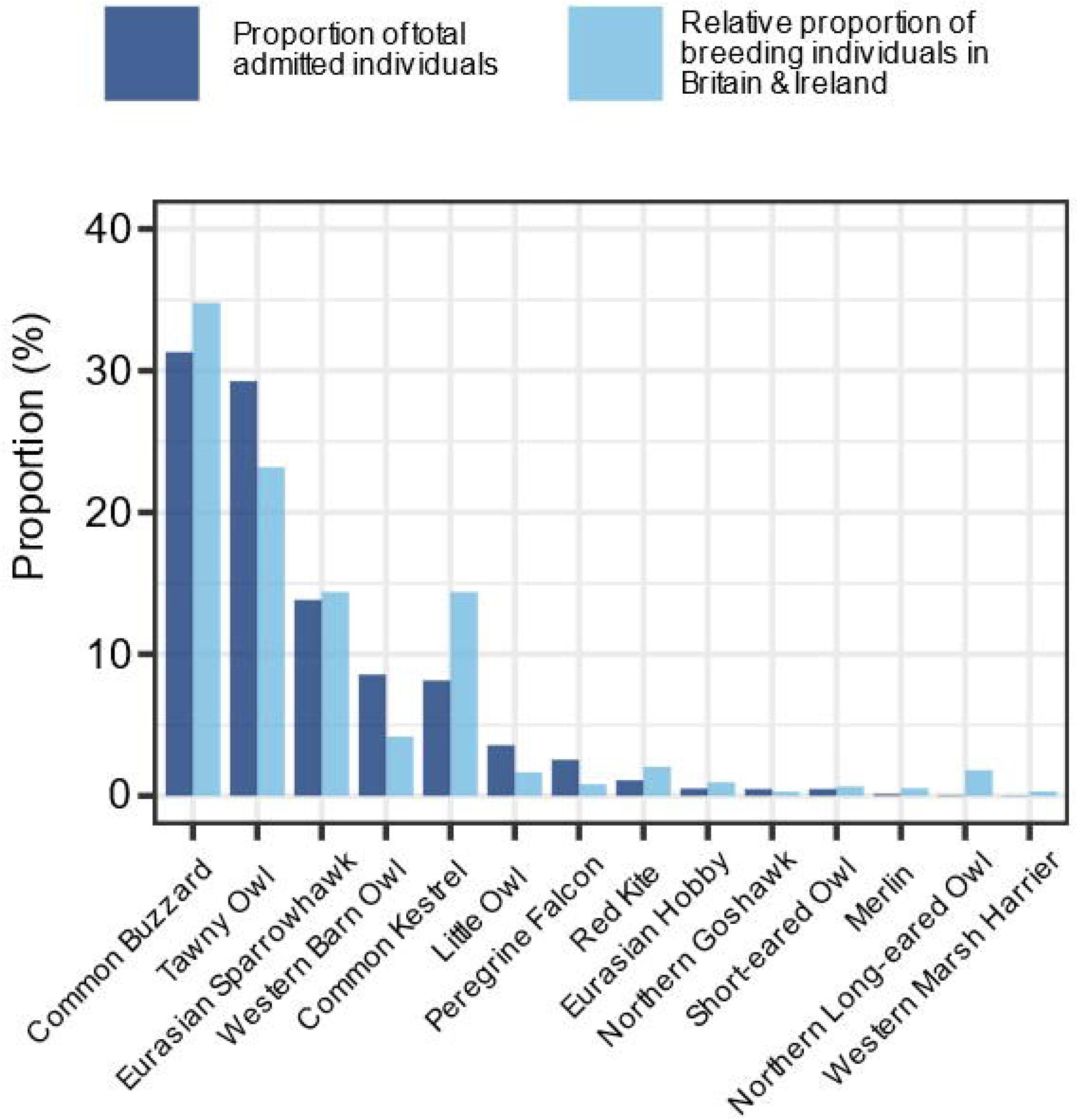
Proportion of total number of admitted individual raptors to four wildlife rehabilitation centres in England and Wales between 2001-2019, compared to the relative proportion of breeding individuals, per species, occurring in Britain and Ireland (data extracted from the BTO BirdFacts database https://www.bto.org/understanding-birds/birdfacts; Robinson 2005).

## DISCUSSION

This study examines, over time, causes of morbidity and mortality for 14 raptors admitted to four wildlife rehabilitation centres in England and Wales, and explores how urbanisation affects causes of admission.

Similar to other studies, unknown trauma accounted for most raptor admissions to wildlife rehabilitation centres (WRC) (see Wendell *et al*. 2002; Rodríguez *et al*. 2010; Mariacher *et al*. 2016; Smith *et al*. 2018; Garcês *et al*. 2019). For example, Molina-López *et al*. (2011) found that trauma accounted for 50% of raptor admissions to a WRC in Spain, with the cause of injury unascertainable for more than half of these. Trauma admissions were also most numerous (56%) in a study of 3,212 raptor admissions to a WRC in New York State, USA (Hanson *et al*. 2021). In South Africa, analysis of eight years of admissions data for 39 raptor species revealed that vehicle and building collisions were the most common cause of admission (Thompson *et al*. 2013), and another South African study found that 52% of all admissions for 33 raptor species were also due to collision-related injuries (Maphalala *et al*. 2021). In our study, collision trauma (both building and vehicle collisions) comprised 56% of all identified admissions and a third of all admissions. In contrast, a 10-year study conducted in Gran Canaria found that 65% of raptor admissions were non-trauma related, e.g., orphaned young birds, with trauma amounting to only around 35% of total admissions (Montesdeoca *et al*. 2017b).

Predominate causes of admission to WRC may vary by country. In Jordan illegal possession and the transport of raptors was the most common admission cause to a single WRC centre between 2017-2018, with trauma cases being the second most frequent admission cause (Al Zoubi *et al*. 2020). A recent study from the Czech Republic reported more than a third of all admissions of 12,923 Common Kestrels to 34 rehabilitation centres were due to nestlings/orphans (Lukesova *et al*. 2021). In our study orphans accounted for 14% of total kestrel admissions, and together with vehicle collisions were the most frequent admission cause for this species.

In our study, nearly 60% of admitted birds either died or were euthanised. Admissions due to anthropogenic causes had a higher mortality rate (57%) than natural causes (40%), and our more refined analysis suggested that infection/parasite admissions were associated with the highest mortality rates (90%), whereas orphaned birds were associated with the lowest mortality rate (16%). Raptors admitted due to being orphaned tend to have higher survival probabilities than those admitted for other reasons, as evidenced by existing studies (Hanson *et al*. 2021; Lukesova *et al*. 2021 [see ‘Nestlings’ and ‘Incubation’ in Table 3]).

### Influence of urbanisation on identified causes of admission

Level of urbanisation was significantly associated with certain admission causes, with building collisions, persecution and unknown trauma admissions more likely to occur in more urbanised areas, but with vehicle collisions more likely in rural areas. Compared to diurnal species, nocturnal species are more susceptible to blinding by vehicle headlights (Bullock *et al*. 2011; Thompson *et al*. 2013). Collisions between Tawny Owls and vehicles have been shown to be more common on roads surrounded by increased tree density (Gomes *et al*. 2009) where connectivity between territories is higher (Santos *et al*. 2013; Gagné *et al*. 2015), i.e., more rural areas, and may explain why vehicle collisions were the most frequent identified admission cause for Tawny Owls in our study. Common Buzzards were the most numerous diurnal species hit by vehicles; the species is less able to adapt to urban habitats (Palomino & Carrascal 2007) and is also a frequent scavenger of roadkill carcasses in rural areas (Young *et al*. 2014; Schwartz *et al*. 2018), which may further explain the increase in vehicle collisions in more rural areas. Vehicle collisions were also the most common admission cause for Western Barn Owls totalling 40% of admissions and were also the most likely cause of death for the species in another study conducted in Britain between 1963-1996 (Newton *et al*. 1997).

Building collisions were more likely to occur in urban areas with the Eurasian Sparrowhawk being the most frequent species admitted for this reason. This species is an urban adapter often breeding in these environments (Thornton *et al*. 2017) employing a high-speed attack strategy when hunting avian prey (Newton 1986). Important causes of mortality have been attributed to collision-based trauma particularly with windows (Newton *et al*. 1999). A study by Kelly & Bland (2006) analysed 202 admissions of Eurasian Sparrowhawk to a WRC in England, 32% of admissions were due to collisions, i.e., vehicle and building/window collisions, which is an identical percentage to our findings for this species admitted to four WRC, suggesting that collision-based injuries (and/or death) are relatively common for the species in England and Wales (Newton *et al*. 1999).

Recently, Crespo *et al*. (2021) found a positive relationship between the number of human inhabitants and avian gunshot admissions in the Valencian region of Spain, the majority of casualties being raptors. We did not explore the effects of human population densities on admission causes, however, we found that persecution admissions (i.e., gunshots, poisoning and traps/snares) increased in urban areas. Assuming that human population densities correlate with urban land cover, our results are in line with those of Crespo *et al*. (2021). Despite this, in Britain it is well-documented that human-raptor conflict often occurs in rural areas such as grouse moors (Thirgood *et al*. 2001; Melling *et al*. 2018; Murgatroyd *et al*. 2019; Newton 2021), although there is no active grouse moor management within our study area, and this pattern might well change if these issues were explored at a larger scale incorporating a wider range of habitat types.

The lack of randomisation (Molina-López *et al*. 2011), restricted geographic study area and small sample sizes for less abundant species (e.g., a single admission for the Western Marsh Harrier and no admissions of species such as the Hen Harrier (*Circus cyaneus*) despite overlap with the species’ distribution in Wales) further limit our ability to explore trends in causes of injury and death for all raptor species occurring throughout England and Wales. Peregrine Falcons were over-represented in our admissions data by 103%. This may be due to recent estimates suggesting that the species’ population size has increased in lowland parts of England along with the overall UK population (Wilson *et al*. 2018), and/or may be due to the species’ well-known use of urban habitats (Kettel *et al*. 2019) subsequently increasing the chance of members of the public encountering injured falcons. Conversely, Northern Long-eared Owls were under-represented in our admissions data by 187%, totalling just two admissions over the study period. In Britain & Ireland, the species’ estimated breeding population size *(ca.* 7800 individuals) is larger than that of other species that were more numerous within our admissions data, e.g., Little Owl (118 admissions; *ca*. 7200 breeding individuals), Northern Goshawk (16 admissions; *ca*. 1240 breeding individuals) and Peregrine Falcon (84 admissions; *ca*. 3500 breeding individuals). Northern Long-eared Owls are nocturnally active and secretive (Petty *et al*. 2003), preferring to use habitats away from human disturbance (Martínez & Zuberogoitia 2004), which may partially explain the low numbers observed in our data.

Admission cause in most cases was based upon details from the finder of the bird (usually a member of the public) and initial assessment by a trained wildlife carer. A veterinary professional (veterinary surgeon or registered veterinary nurse) was usually not involved at this stage, so a definitive clinical diagnosis was not made. The centres involved however, all have very experienced and well-trained staff, with an ability to make a good initial assessment of the bird. However, identification accuracy between WRC and trained wildlife carers is unlikely to be equal, which should be considered when making inferences from these data.

For 77% of admissions sex was not determined, constraining our ability to compare admission causes between the sexes. However, the majority of admitted birds were able to be assigned to a broad age category allowing for age-related demographic comparisons. Nevertheless, 60% of admissions were of adult birds which support results from WRC in the USA (Hernandez *et al*. 2018) and Greece (Komnenou *et al*. 2005). The remaining 40% of admissions comprised juvenile birds and similar patterns have been observed elsewhere; for example, 42% of Northern Long-eared Owl admissions (Italy; Mariacher *et al*. 2016) and 32% of all raptor admissions (Spain; Molina-López *et al*. 2011) being juveniles.

Relative to anthropogenic admissions, natural admissions are likely to be under-represented in our data due to the majority going unreported (Real *et al*. 2001; Newton 2002). The reliance of reports from members of the public means that there is a likely bias towards anthropogenic admission causes. Building and vehicle collisions are more likely to be reported by members of the public by chance than persecution, i.e., illegal activities such as poisoning, gunshot and trap/snare events. Our data may also include a survivability bias with members of the public more likely to report injured birds that are still alive than those that have already died, inhibiting reliable injury and death estimates at local raptor population-levels.

Alternative monitoring methods such as satellite telemetry are more reliable sources for capturing illegal wildlife crimes, as demonstrated by Murgatroyd *et al*. (2019) who examined patterns of Hen Harrier disappearances over grouse moors in northern England as a result of suspected illegal killing. In addition, Panter *et al*. (2021) used satellite telemetry to estimate survival in wintering Red Kites in south-western Europe and Oppel *et al*. (2021) coupled satellite telemetry and on-the-ground surveys to explore Egyptian Vulture *(Neophron percnopterus*) mortalities along their migratory routes. However, using such technology is often costly and requires specialist skill. Analysis of admissions data is cost-effective and requires little investment other than time, and many WRC often keep records of wildlife admissions for their own purposes as demonstrated in this study.

### Implications

Admissions data from WRC have the potential to form important baseline data guiding conservation activities. For example, gunshot admissions data from Greece has been used to advise governmental agencies responsible for hunting regulations (Mazaris *et al*. 2008) and seasonal cumulative indices have been calculated to explore the potential ecological impacts on local raptor populations in Spain (Molina-López *et al*. 2011). Some 39% of raptors were released back into the wild following treatment, however, release does not equate to successful reintroduction back into breeding populations. Post-release monitoring of individuals, for example via identification of individuals using leg bands and coupled with field surveys, is strongly encouraged. This provides additional conservation value to admissions data and also allows for post-release welfare checks to be made on the bird.

Building and vehicle collisions posed the highest identified risk to raptors in our study area. Increased traffic densities and vehicle speeds have been shown to increase bird-vehicle collision mortalities (Erritzoe *et al*. 2003). Identification of vehicle collision hotspots along road networks is recommended and predictive modeling has been applied at the landscape- and local-scale to improve road safety (Malo *et al*. 2004). Window decals have successfully reduced average monthly bird-window collisions by 84% (Ocampo-Peñuela *et al*. 2016). Application of collision prevention decals to the exterior surface of windows (Klem & Saenger 2012), or tinting of windows (Erickson *et al*. 2005), are viable solutions to prevent bird-building collisions and citizen science can assist with community-level implementations.

Transformation of natural habitats into human-modified environments has been shown to negatively affect raptor communities, resulting in lower abundances, species richness and diversity (Carette *et al*. 2009). Despite this, some raptor species have shown resilience and even proliferation of urban environments (Cooke *et al*. 2018; Kettel *et al*. 2019; Panter *et al*. 2020). For example, Sumasgutner *et al*. (2020) found that breeding Peregrine Falcon pairs were more likely to breed and bred earlier in more urbanised areas, compared to their more rural conspecifics, but breeding success may be compromised in more urban areas for some species, e.g., Common Kestrels (Kettel *et al*. 2018).

Many threats persist for raptors in England and Wales, however, have not changed substantially over the past two decades. Our findings provide baseline data on the causes of morbidity and mortality of raptors throughout our study area. Threats associated with urban areas, such as building collisions, may increase over time in line with human population growth and subsequent urban expansion. There is potential for future studies to build on our results in an applied context, for example, investigating the financial costs of vehicle damage as a result of vehicle-wildlife collisions.

## ACKNOWLEDGEMENTS

We would like to thank Marlies Hebdon (Secret World Rescue) for assistance with data collection and D. Zanders for her useful comments on the draft manuscript. We would like to further thank the following photographers for use of their Creative Commons images: Mark Kilner (CC BY-NC-SA 2.0 Flickr), YvonneHuijbens (CC0 Pixabay), Bых Πыхманн (CC BY-SA 3.0 Wikimedia Commons), Tambako The Jaguar (CC BY-ND 2.0 Flickr), Imran Shah (CC BY-SA 2.0 Flickr), Peter K. Burain (CC BY 4.0) and Sue Cro (CC BY-NC 2.0 Flickr).

## APPENDICES

### Appendix A

Admission type, cause, code and descriptions for 3305 admission records of raptors admitted to four wildlife rehabilitation centres in the United Kingdom between 2001-2019.

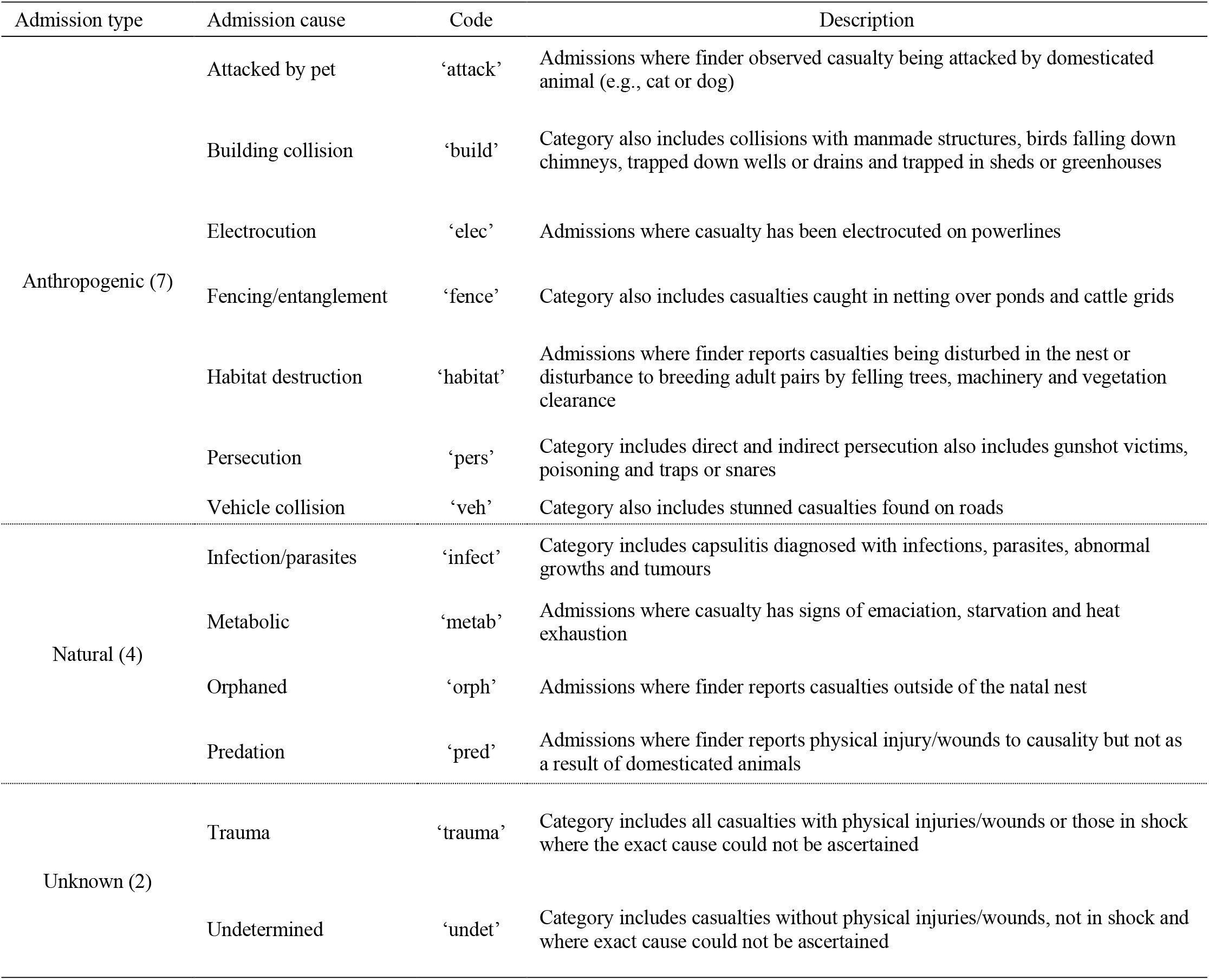

### Appendix B

An overview of models used to explore trends over time and effects of urbanisation on raptor admissions to four wildlife rehabilitation centres (WRC) in England and Wales between 2001-2019. GLM = Generalized Linear Model; GLMM = Generalized Linear Mixed Model; GBH = Gower Bird Hospital only.

| Model type | Terms |  |  | Error distribution | Link function | Data set | Data included (admission types) |  |  |
| --- | --- | --- | --- | --- | --- | --- | --- | --- | --- |
|  | Response | Explanatory | Random |  |  |  | Anthropogenic | Natural | Unknown |
| GLM | fate (binary) | admission type/cause | - | binomial | logit | All WRC | ✓ | ✓ |  |
| GLM | total admission counts | year | - | quasi-Poisson | - | GBH | ✓ | ✓ | ✓ |
| cbind GLM | admission counts per type (anthropogenic vs. natural) | year | - | binomial | - | GBH | ✓ | ✓ |  |
| GLM | admission counts per species | year | - | quasi-Poisson | - | GBH | ✓ | ✓ | ✓ |
| GLM | relative proportion of admission counts by per cause | year | - | quasi-Poisson | - | GBH | ✓ | ✓ | ✓ |
| GLMM | anthropogenic (binary) | % urban land cover | centre ID | binomial | logit | All WRC | ✓ |  |  |
| GLMM | natural (binary) | % urban land cover | centre ID | binomial | logit | All WRC |  | ✓ |  |
| GLMM | unknown (binary) | % urban land cover | centre ID | binomial | logit | All WRC |  |  | ✓ |
| GLMM | cause (binary) | % urban land cover | centre ID | binomial | logit | All WRC | ✓ | ✓ | ✓ |

### Appendix C

Pairwise comparisons of mortality probabilities for raptors admitted to four wildlife rehabilitation centres in England and Wales between 2001-2019, presented by identified admission types (above dashed line) and admission causes (below dashed lines). Analyses conducted using a series of Generalized Linear Models with binomial error distributions and ‘logit’ link functions. **Bold** = statistically significant comparisons, SE = standard error.

| Contrast | Estimate ± SE | *z* | *P* |
| --- | --- | --- | --- |
| **anthropogenic - natural** | **0.668 ± 0.103** | **6.483** | **< 0.0001** |
| attacked - building collision | 0.509 ± 0.369 | 1.385 | 0.904 |
| attacked - fencing/entanglement | 0.657 ± 0.422 | 1.556 | 0.828 |
| **attacked - infection/parasites** | **-1.601 ± 0.494** | **-3.241** | **< 0.05** |
| attacked - metabolic | 0.250 ± 0.386 | 0.650 | 0.999 |
| **attacked - orphaned** | **2.238 ± 0.380** | **5.895** | **< 0.0001** |
| attacked - persecution | 0.490 ± 0.417 | 1.176 | 0.962 |
| attacked - predation | -0.076 ± 0.509 | -0.150 | 1.000 |
| attacked - vehicle collision | 0.171 ± 0.355 | 0.484 | 1.000 |
| building collision - fencing/entanglement | 0.147 ± 0.271 | 0.546 | 1.000 |
| **building collision - infection/parasites** | **-2.116 ± 0.373** | **-5.665** | **< 0.0001** |
| building collision - metabolic | -0.259 ± 0.208 | -1.244 | 0.947 |
| **building collision - orphaned** | **1.728 ± 0.197** | **8.759** | **< 0.0001** |
| building collision - persecution | -0.019 ± 0.262 | -0.074 | 1.000 |
| building collision - predation | -0.586 ± 0.392 | -1.495 | 0.858 |
| building collision - vehicle collision | -0.338 ± 0.144 | -2.341 | 0.317 |
| **fencing/entanglement - infection/parasites** | **-2.259 ± 0.426** | **-5.298** | **< 0.0001** |
| fencing/entanglement - metabolic | -0.407 ± 0.294 | -1.385 | 0.904 |
| **fencing/entanglement - orphaned** | **1.580 ± 0.286** | **5.525** | **< 0.0001** |
| fencing/entanglement - persecution | -0.167 ± 0.334 | -0.500 | 1.000 |
| fencing/entanglement - predation | -0.733 ± 0.443 | -1.655 | 0.774 |
| fencing/entanglement - vehicle collision | -0.485 ± 0.253 | -1.923 | 0.597 |
| **infection/parasites - metabolic** | **1.852 ± 0.390** | **4.750** | **< 0.0001** |
| **infection/parasites - orphaned** | **3.839 ± 0.384** | **9.997** | **< 0.0001** |
| **infection/parasites - persecution** | **2.092 ± 0.421** | **4.971** | **< 0.0001** |
| infection/parasites - predation | 1.525 ± 0.512 | 2.979 | 0.071 |
| **infection/parasites - vehicle collision** | **1.773 ± 0.360** | **4.930** | **< 0.0001** |
| **metabolic - orphaned** | **1.987 ± 0.228** | **8.710** | **< 0.0001** |
| metabolic - persecution | 0.239 ± 0.286 | 0.839 | 0.996 |
| metabolic - predation | -0.326 ± 0.408 | -0.800 | 0.997 |
| metabolic - vehicle collision | -0.078 ± 0.184 | -0.427 | 1.000 |
| **orphaned - persecution** | **-1.747 ± 0.278** | **-6.291** | **< 0.0001** |
| **orphaned - predation** | **-2.314 ± 0.403** | **-5.745** | **< 0.0001** |
| **orphaned - vehicle collision** | **-2.066 ± 0.172** | **-12.039** | **< 0.0001** |
| persecution - predation | -0.566 ± 0.438 | -1.293 | 0.934 |
| persecution - vehicle collision | -0.318 ± 0.243 | -1.310 | 0.929 |
| predation - vehicle collision | 0.248 ± 0.380 | 0.653 | 0.999 |

### Appendix D

Parameter estimates from the Generalized Linear Models, fitted with a quasi-Poisson error distribution to account for overdispersion, examining trends over time for the seven most common species admitted to Gower Bird Hospital between 2001-2019. **Bold** = statistically significant causes. N = number of admissions, SE = standard error, df = degrees of freedom.

| Species | N | Estimate ± SE | *t* | df | *P* |
| --- | --- | --- | --- | --- | --- |
| **Common Buzzard (*Buteo buteo*)** | **470** | **-0.034 ± 0.014** | **-2.407** | **17** | **< 0.05** |
| Tawny Owl (*Strix aluco*) | 323 | -0.021 ± 0.019 | -1.112 | - | 0.282 |
| Eurasian Sparrowhawk (*Accipiter nisus*) | 193 | -0.008 ± 0.016 | -0.521 | - | 0.609 |
| Western Barn Owl (*Tyto alba*) | 105 | -0.003 ± 0.029 | -0.117 | - | 0.908 |
| **Common Kestrel (*Falco tinnunculus*)** | **77** | **-0.081 ± 0.020** | **-4.031** | **-** | **< 0.001** |
| Peregrine Falcon (*Falco peregrinus*) | 44 | 0.028 ± 0.030 | 0.926 | - | 0.368 |
| **Red Kite (*Milvus milvus*)** | **34** | **0.155 ± 0.033** | **4.703** | **-** | **< 0.001** |

### Appendix E

Percentage difference between the relative proportion of breeding individuals in Britain & Ireland, and the proportion of individuals, per species, admitted to four wildlife rehabilitation centres in England and Wales between 2001-2019. *Data derived from the British Trust for Ornithology’s BirdFacts database (Robinson 2005; https://www.bto.org/understanding-birds/birdfacts).

| Species | Number of<br>breeding<br>individuals in<br>Britain &<br>Ireland* | Proportion<br>breeding<br>individuals in<br>Britain &<br>Ireland | Number of<br>admitted<br>individuals | Proportion<br>of total<br>admitted<br>individuals | % Difference<br>(breeding<br>individuals<br>vs. admitted<br>individuals) |
| --- | --- | --- | --- | --- | --- |
| Peregrine Falcon (*Falco peregrinus*) | 3500 | 0.8 | 84 | 2.5 | 103.2 |
| Little Owl (*Athene noctua*) | 7200 | 1.7 | 118 | 3.6 | 72.5 |
| Western Barn Owl (*Tyto alba*) | 18000 | 4.2 | 283 | 8.6 | 68.9 |
| Northern Goshawk (*Accipiter gentilis*) | 1240 | 0.3 | 16 | 0.5 | 50.9 |
| Tawny Owl (*Strix aluco*) | 100000 | 23.2 | 967 | 29.3 | 23.1 |
| Eurasian Sparrowhawk (*Accipiter nisus*) | 62000 | 14.4 | 457 | 13.8 | -3.9 |
| Common Buzzard (*Buteo buteo*) | 150000 | 34.8 | 1035 | 31.3 | -10.5 |
| Short-eared Owl (*Asio flammeus*) | 2820 | 0.7 | 16 | 0.5 | -29.9 |
| Common Kestrel (*Falco tinnunculus*) | 62000 | 14.4 | 269 | 8.1 | -55.5 |
| Eurasian Hobby (*Falco subbuteo*) | 4100 | 1.0 | 17 | 0.5 | -59.6 |
| Red Kite (*Milvus milvus*) | 8800 | 2.0 | 36 | 1.1 | -60.8 |
| Merlin (*Falco columbarius*) | 2300 | 0.5 | 4 | 0.1 | -126.0 |
| Western Marsh Harrier (*Circus aeruginosus*) | 1285 | 0.3 | 1 | 0.0 | -163.1 |
| Northern Long-eared Owl (*Asio otus*) | 7800 | 1.8 | 2 | 0.1 | -187.1 |

